# Development of a deep amplicon sequencing method to determine the proportional species composition of piroplasm haemoprotozoa as an aid in their control

**DOI:** 10.1101/580183

**Authors:** Umer Chaudhry, Qasim Ali, Imran Rashid, Muhammad Zubair Shabbir, Muhammad Abbas, Muhammad Numan, Mike Evans, Kamran Ashraf, Ivan Morrison, Liam Morrison, Neil D. Sargison

**Author notes:** Corresponding author: Umer Chaudhry, University of Edinburgh, The Roslin Institute, Easter Bush Veterinary Centre, UK, EH25 9RG.

## Abstract

Piroplasmosis is caused by tick-borne haemoprotozoa of the genera *Theileria* and *Babesia*. These parasitic infections can cause serious impact on the health of livestock and production. Multiple piroplasm species can infect a single host, but reliable molecular diagnostic tools are needed with which to understand the composition of these complex parasite communities. *Theileria* and *Babesia* vary in their epidemiology, drug sensitivity, pathogenicity and interaction of co-infecting species, but are similar in the animals, become persistent carriers after recovery from primary infection, acting as reservoir hosts. Here, we describe for the first time the use of a deep amplicon sequencing platform to identify proportions of piroplasm species in co-infecting communities and develop the concept of a “haemoprotobiome”. First, four phenotypically-verified species of *Theileria* and *Babesia* were used to prepare mock pools with random amounts of the parasites and amplified with four different numbers of PCR cycles to assess sequence representation of each species. Second, we evaluated the detection threshold of the deep amplicon sequencing assay for each of the four species and to assess the accuracy of proportional quantification of all four species. Finally, we applied the assay to the field samples to afford insight of the species composition of piroplasm communities in small and large ruminants in the Punjab province of Pakistan. The “haemoprotobiome” concept has several potential applications in veterinary and human research, including understanding of responses to drug treatment; parasite epidemiology and ecology; species interactions during mixed infections; and parasite control strategies.

## Introduction

Haemoprotozoa are ubiquitous and amongst the most successful organisms in the biosphere, possessing an incredible ability to adapt too many different environments and niches (Nene *et al.*, 2016). Among these, *Theileria* and *Babesia* are arguably the most economically important and highly pathogenic neglected livestock parasites, causing theileriosis and babesiosis. Different species of *Theileria* and *Babesia* occur globally, impacting on the health, welfare and production of livestock (Jabbar *et al.*, 2015). It is estimated that about approximately 80% of the world’s cattle population is infected, causing economic loss due to high morbidity and mortality and threatening food security in many livestock dependent communities (Sivakumar *et al.*, 2014). Theileriosis and babesiosis are particularly important in subtropical regions, where efficient ruminant livestock production is critical to the wellbeing and poverty alleviation of smallholder farmers (Gubbels *et al.*, 1999).

Next generation genomic resources have potential applications in the diagnosis, surveillance, treatment and control of haemoprotozoan diseases, as well as in the evaluation of parasite population responses to drug treatments and other control strategies. Determination of sequence variations in the hyper-variable 18S rDNA cistron can discriminate between haemoprotozoan parasites (Gubbels *et al.*, 1999) and overcome limitations of traditional gross parasitological methods for the diagnosis of haemoprotozoa at species level (Agudelo *et al.*, 2013; Haanshuus *et al.*, 2013; Lee *et al.*, 2015; Lefterova *et al.*, 2015; Mens *et al.*, 2006; Rougemont *et al.*, 2004; Steenkeste *et al.*, 2009). Various PCR methods (reverse line blot (RLB)-PCR, quantitative (q)PCR, multiple PCR) have been described to amplify the 18S region for sequence determination, but these are low throughput, hence relatively expensive, and can be error-prone (Bilgic *et al.*, 2013; Chaisi *et al.*, 2013; Gubbels *et al.*, 1999; Kundave *et al.*, 2018). These methods depend on the use of species-specific primers and probes, hence can only identify the tested species. In contrast, high throughput deep amplicon sequencing using the Illumina Mi-Seq platform is relatively low-cost and involves fewer PCR cycles, hence is potentially less error prone. The method has transformed the study of bacterial (microbiome) (Gloor *et al.*, 2010; Rogers & Bruce, 2010) and nematode parasites (nemabiome) (Avramenko *et al.*, 2015b; Chaudhry & Sargison, 2018), opens new areas of research in the study of protozoan parasite communities. The method has the potential to accurately identify and provide relative quantification of co-infecting species of haemoprotozoa. The use of primers binding to conserved sites and analysis of up to 600 bp sequence reads allows previously unanticipated protozoan species to be detected. The use of barcoded primers allows a large number of samples to be pooled and sequenced in a single Mi-Seq run, making the technology suitable for high throughput analysis (Avramenko *et al.*, 2015b). By multiplexing the barcoded primer combinations, it is possible to run 384 samples at once on a single Illumina Mi-Seq flow cell, helping to reduce the cost.

Here we report the development of a method of deep amplicon sequencing of the hyper-variable 18S rDNA region using Illumina Mi-seq platform and the validation of its accuracy for species quantification of *Theileria* and *Babesia* communities present in field samples.

## Materials and methods

### Parasite material and genomic DNA preparation

*Theileria* and *Babesia* species of large and small ruminants were chosen in this study; first, because the availability of phenotypically verified isolates allowed the creation of ‘mock pools’ of the parasites to assess the sequence representation and the proportions of each of species and second, because the availability of piroplasm positive buffalo, cattle and sheep blood samples allowed us to demonstrate the method in the field. Five phenotypically verified piroplasm isolates (*Theileria annulata, Theileria parva, Theileria lestoquardi, Babesia bigemina and Babesia bovis*) had been derived from different parts of the world and were held in common reference cell lines at the University of Edinburgh, Roslin Institute, UK.

Mock pools of the four isolates (*T. annulata, T. parva, B. bigemina* and *B. bovis*) were created, by taking 50 µl aliquots from each of the species. Three replicates from the four mock pools were made and amplified using four different numbers of PCR cycles (25X, 30X, 35X and 40X), to assess the sequence representation of each species. Three replicates of the nine mock pools were used to test the detection threshold of the deep amplicon sequencing method and to show the proportions of each of the species present. The description of the nine separate mock pools are following: [Mix1 (*T. annulata, T. parva, B. bigemina* and *B. bovis*), Mix2 (*T. annulata, T. parva* and *B. bovis*), Mix3 (*T. annulata, T. parva* and *B. bigemina*), Mix4 (*T. parva, B. bigemina* and *B. bovis*), Mix5 (*T. annulata, B. bigemina* and *B. bovis*), Mix6 (*B. bovis*), Mix7 (*B. bigemina*), Mix8 (*T. annulata*), Mix9 (*T. parva*)]. Sample series were prepared with each of the main species being absent from one mixture (Fig. 3). gDNA was prepared using 1,000 μl Direct PCR lysis reagent (Viagen), 50 μl proteinase K solution (Qiagen), and 50μl 1M dithiothreitol (DTT). To extract gDNA, 50 μl of aliquots from each of the species was transferred into a fresh tube and centrifuged for 5 minutes, before removing the supernatant and mixing with 25 μl lysis buffer (Viagen) and Proteinase K (New England BioLabs). Lysates were prepared using standard methods (Chaudhry *et al.*, 2015a).

A total of 79 piroplasm positive buffalo blood samples, 86 piroplasm positive cattle blood samples and 18 piroplasm positive sheep blood samples were collected from veterinary clinics throughout the Punjab province of Pakistan between 2017 and 2018. The procedures involved jugular venipuncture and withdrawal of 5 ml of intravenous blood into an EDTA tubes, followed by storage at −20 □C. Discussions were held with key administrative and community leaders to raise awareness of the study, and encourage households to participate. Samples were taken by trained para-veterinary workers under the supervision of local veterinary staff. The study was approved by the Institutional Review Board of the University of Veterinary and Animal Sciences Punjab, Pakistan. Thick and thin blood smears were stained with 10% Giemsa solution and examined under the oil immersion (100 x) lens to identify piroplasms. To extract gDNA, 50 μl of blood from each of the piroplasm positive samples was used as template, and the DNA was extracted according to the protocols described in the TIANamp blood DNA kit (Beijing Biopeony Co. Ltd).

### Deep amplicon sequencing

The overall scheme of the deep amplicon sequencing approach using Illumina MiSeq platform has been described below and shown in S1 Fig.

#### Adapter PCR amplification of 18S rDNA locus

A 463 to 504 bp fragments encompassing parts of the 18S rDNA spanning V4 hyper-variable region of haemoprotozoa was used for 1^st^ round adapter PCR amplification. The primers were altered from the normal primers set (RLB_For, RLB_Rev) previously described by Gubbels *et al.* (1999). The adapters were added to each primer to allow the successive annealing and N is the random number of nucleotides included between the adapter and primer set (S1 Table). Equal proportions of four forward (RLB_For, RLB_For-1N, RLB_For-2N, RLB_For-3N) and four reverse primers (RLB_Rev, RLB_Rev-1N, RLB_Rev-2N, RLB_Rev-3N) were mixed and used for the adapter PCR with following conditions: 10 uM forward and reverse adapter primer, 10 mM dNTPs, 0.5 U DNA polymerase enzyme, 5X buffer (KAPA Biosystems, USA) and 1 μl of gDNA. The thermocycling conditions were 95°C for 2 minutes, followed by 35 cycles of 98°C for 20 seconds, 60°C for 15 seconds, 72°C for 15 seconds and a final extension of 72°C for 5 minutes. The PCR product was purified with AMPure XP Magnetic Beads (1X) using a special magnetic stand and plate according to the protocols described by Beckman coulter, Inc.

#### Barcoded PCR amplification of 18S rDNA locus

The 2^nd^ round barcoded PCR was performed by using sixteen forward and twenty-four reverse barcoded primers obtained from Illumina MiSeq protocols (S2 Table). Repetitions of the forward and reverse barcoded primers in different samples were avoided. The barcocded PCR was performed with the following conditions: 10 uM barcoded forward (N501 to N508, N510 to N517) and reverse (N701 to N712 and N713 to N724) primers, 10 mM dNTPs, 0.5 U DNA polymerase enzyme, 5X buffer (KAPA Biosystems, USA) and 2 μl of adaptor PCR product as DNA template. The thermocycling conditions were 98°C for 45 seconds, followed by 7 cycles of 98°C for 20 seconds, 63°C for 20 seconds, and 72°C for 2 minutes. The PCR product was purified with AMPure XP Magnetic Beads (1X) according to the protocols described by Beckman coulter, Inc.

#### Illumina Mi-Seq run, data handling and bioinformatics analysis

The pooled library was prepared with 10 μl of barcoded PCR bead purified product from each individual sample and measured with KAPA qPCR library quantification kit (KAPA Biosystems, USA), before being run on an Illumina Mi-Seq Sequencer using a 600-cycle pair end reagent kit (Mi-Seq Reagent Kits v2, MS-103-2003) at a concentration of 15 nM with addition 25% Phix Control v3 (Illumina, FC-11-2003). The Mi-Seq separates all sequencing data by sample during post-run processing using recognised barcoded indices to generate FASTAQ files. The data was analysed with our own pipeline. Briefly, the consensus sequences taxonomy library of the 18S rDNA of *Theleria* and *Babesia* species has been described in the next section of the materials and methods. Illumina Mi-seq data analysis was performed using Mothur v1.39.5 software (Schloss *et al.*, 2009) and the Illumina Mi-seq SOPs (Kozich *et al.*, 2013). Overall, thousands of 18S rDNA reads were generated from the data set of five phenotypically verified *Theileria* and *Babesia* isolates and piroplasm positive field samples [buffalo (n=79), cattle (n=86) and sheep (n=18)]. In summary, raw paired reads were made into contigs, and those were too long, or ambiguous bases were removed. The sequence data were trimmed near the region of forward and reverse primers and aligned with the consensus sequence library. The sequence data that did not match with the 18S rDNA consensus sequences taxonomy library was discarded as a trace amplicon contamination. One challenge is to detect the trace amplicon contamination that may occur during sample handling subsequent to the adapter and barcoded PCR amplification. However, minor sequence contamination does not affect the interpretation of the relative species proportions, since the trace amplicon contamination was always less than the number of sequences generated from the samples. Consequently, contaminated sequences can be distinguished from genuine data. Further, the very small changes in species proportions associated with trace contamination do not change the biological interpretation of the data.

### Consensus sequence libraries preparation of phenotypically verified Theileria and Babesia species

Deep amplicon sequencing of the 18S rDNA of three *Theileria* species (*T. annulata, T. parva, T. lestoquardi*), and two *Babesia* species (*B. bigemina and B. bovis*) was described in the previous section. The generated reads were compared to the 18S sequences of other haemoprotozoan species published in NCBI GeneBank to account for any additional genetic diversity. We obtained the GeneBank sequences of nine *Theileria* species (*T. orientalis, T. sergenti, T. buffeli, T. velifera, T. taurotragi, T. mutant, T. uilenbergi, T. luwenshuni, T. ovis*) and five *Babesia* species (*B. orientalis, B. occultans, B. ovata, B. major, B. ovis*). The 18S rDNA sequences of twelve *Theleria* and seven *Babesia* species were first aligned in Geneious software and then imported into the CD-HIT software to calculate the number of consensus sequences generated from each species. Finally, 86 consensus sequences were generated from *Theileria* and *Babesia* species of large ruminants, 21 consensus sequences were generated from *Theileria* and *Babesia* species of small ruminants and used as the taxonomy library for data analysis in the previous section. A phylogenetic tree of the 18S rDNA consensus sequences were then constructed using Maximum Likelihood method in MEGA5 software previously described by Chaudhry *et al.* (2015b).

## Results

### Assessment of 18S rDNA genetic diversity of phenotypically verified Theileria and Babesia species

Total 43 consensus sequences of the 18S rDNA locus were identified among eight large ruminant *Theleria* species (*T. annulata=12, T. parva=6, T. orientalis=5, T. sergenti=2, T. buffeli=5, T. velifera=4, T. taurotragi=2, T. mutant=7*) and 43 consensus sequences were identified among six large ruminant *Babesia* species (*B. bigemina=17, B. bovis=15, B. orientalis=2, B. occultans=4, B. ovata=3, B. major=2*) (Fig. 1A). A phylogenetic tree of the 86 consensus sequences demonstrates a distinct clustering between species (Fig 1A). Comparison of the genetic distance of 18S rDNA revealed 87% to 98% homology within the eight *Theileria* species and varied from 77% to 97% homology within the six *Babesia* species. The genetic distance also revealed differences in homology ranging from 72% to 86% between *Theileria* and *Babesia* genera (S3 Table). The most closely related species of *Theileria* and *Babesia* could still be a reliably differentiated by virtue of 18S rDNA sequence variations (S4 Table).

**Fig. 1.**
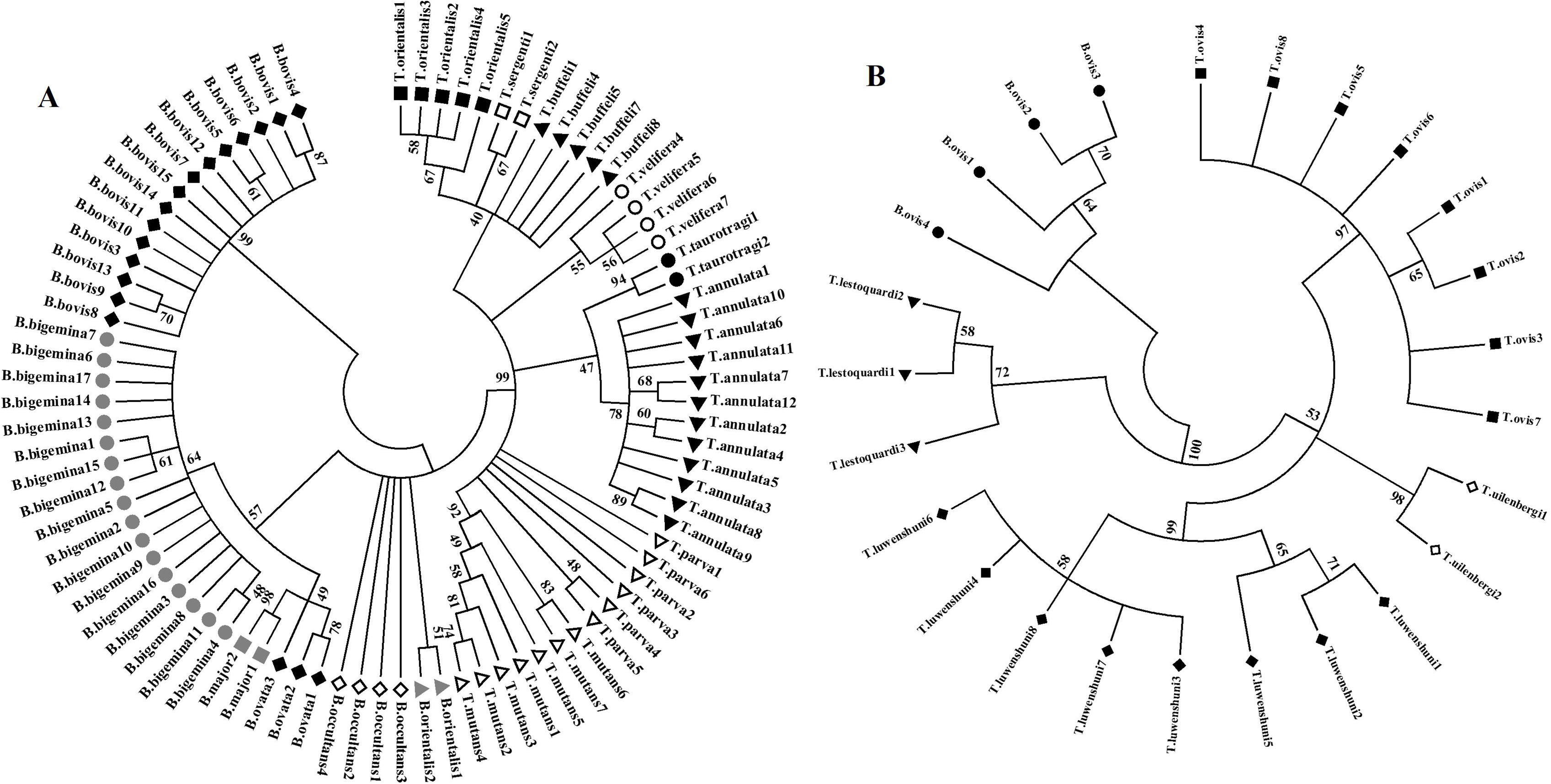
Maximum Likelihood tree of 86 and 21 consensus sequences obtained from the *Theleria* and *Babesia* species. The neighbor-joining tree (Kimura 2-parameter model) was computed with 1000 bootstrap replicates using MEGA5 software created by Biomatters. Each species was identified with different shades bars.

Total 21 consensus sequences of the 18S rDNA locus were identified among four small ruminant *Theleria* species (*T. lestoquardi=3, T. ovis=8, T. uilenbergi=2, T. luwenshuni=8*) and 4 unique haplotypes in *B. ovis* (Fig. 1B). The phylogenetic tree of the 25 consensus sequences showed distinct clustering between species (Fig 1B). Genetic distance comparison of the 18S rDNA revealed 98% to 95% homology within the four *Theileria* species and differences in homology ranging from 81% to 83% between *Theileria* and *Babesia* genus (S3 Table).

### Evaluation of the sequence representation of each of the four species (T. annulata, T. parva, B. bigemina and B. bovis) using different number of PCR cycles

The deep amplicon sequencing assay was performed three independent times from four isolates to assess the sequence representation of each species (Fig 2, S5 Table). The number of cycles for the adapter PCR had no impact on the sequence representation of each species. Each of the species was significantly represented in each number of cycles, based on the number of sequences generated from each mock pool (Fig 2A). Therefore, the effect of the PCR cycle number was statistically analysed by running a Kruskal-Wallis rank sum test for each PCR. There was no statistically substantial differences in the proportion of each species between different numbers of PCR cycles (overall *H*_(3)_<0.04, *p*>0.9). Calculating the mean values based on each cycle of amplification removed further deviation, indicating that level of amplification does not influence the proportional representation of each species. Additionally, it was demonstrated that replicates of the same samples minimize all differences in representation produced by the assay (Fig 2B).

**Fig. 2.**
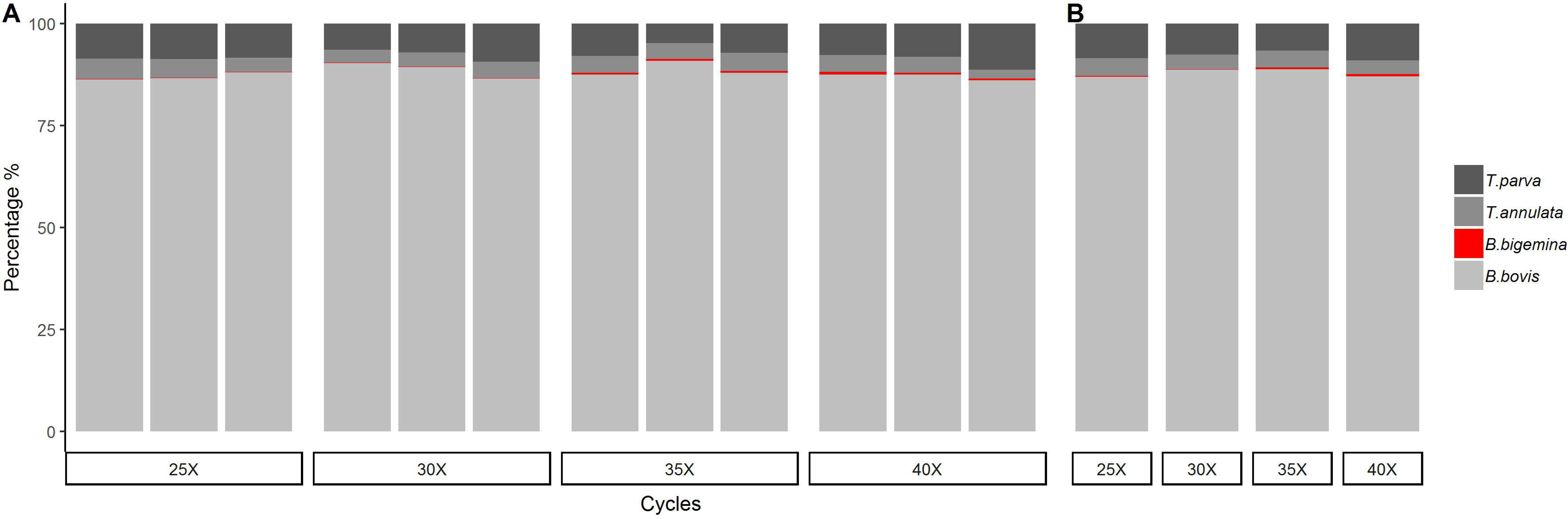
Assessment of species representation bias for the deep amplicon sequencing assay. A gDNA was prepared from a mock pool of the random number of parasites from each of the following species; *T. annulata, T. parva, B. bigemina and B. bovis*. 2A. The sequencing assay was applied three times at 25, 30, 35 and 40 cycles of amplification respectively as denoted on the X-axis and Y-axis shows the percentage proportions of each species. 2B. Replicates were grouped and averaged based on cycles of amplification.

**Fig. 3.**
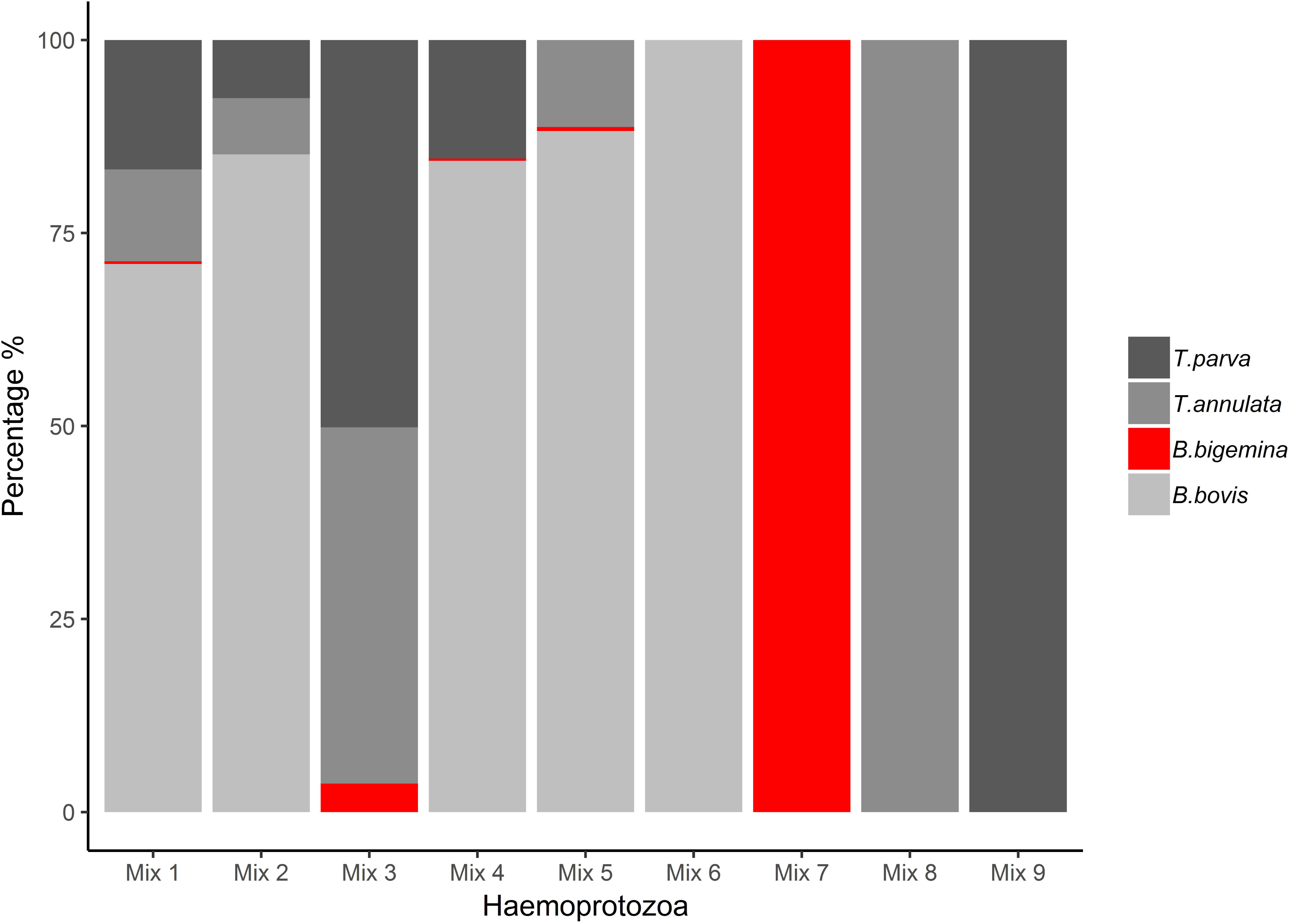
Validation of the deep amplicon sequencing assay. Nine separate mock pools of the random number of parasites from each the four haemoprotozoa species were created. The panel shows the species proportions estimated from the results of the deep amplicon sequencing assay. In each case, the Y-axis shows the percentage proportions of each species and the X-axis indicates the mixing of the mock pools from each species; Mix1(*T. annulata, T. parva, B. bigemina* and *B. bovis*), Mix2(*T. annulata, T. parva* and *B. bovis*), Mix3(*T. annulata, T. parva* and *B. bigemina*), Mix4(*T. parva, B. bigemina* and *B. bovis*), Mix5(*T. annulata, B. bigemina* and *B. bovis*), Mix6(*B. bovis*), Mix7(*B. bigemina*), Mix8(*T. annulata*), Mix9(*T. parva*).

### Validation of the deep amplicon sequencing assay using a mock pools of T. annulata, T. parva, B. bigemina and B. bovis

Three replicates each of known admixtures were created from gDNA from four isolates, to validate the deep amplicon sequencing assay (Fig 3, S6 Table). The mixing of different mocks pools demonstrates the perfect detection threshold of the deep amplicon sequencing method and to show the proportions of each of the species present. The Mix 1 pool contained all four isolates, the results were perfectly matched (Fig 3). The Mix 2 contained *T. annulata, T. parva* and *B. bovis*, Mix 3 contained *T. annulata, T. parva* and *B. bigemina*, Mix 4 contained *T. parva, B. bigemina* and *B. bovis* and Mix 5 contained *T. annulata, B. bigemina* and *B. bovis,* the results were accurate, with no significant variations between replicates (Fig 3). For the pools of 100% of each species (Mix 6, Mix 7, Mix 8, Mix 9), the results are consistently perfect (Fig 3).

### Assessment of the deep amplicon sequencing in the quantification of piroplasm positive blood samples

The deep amplicon sequencing was been applied to the piroplasm positive blood samples of buffalo, cattle and sheep to accurately quantify the levels of haemoprotozoan infection in the field (Fig. 4, S7 Table). Overall, 183 piroplasm positive field samples [buffalo (n=79), cattle (n=86) and sheep (n=18)] were examined to identify *Theileria* and *Babesia* infections. A summary of the results is shown in Fig. 4. In the buffalo and cattle samples, the results demonstrated that the prevalence of *T. annulata* infection was higher than that of *B. bigemina* and *B. bovis* infection. All 79 and 86 samples were *T. annulata* positive, 10/79 buffalo and 10/86 cattle samples were *B. bovis* positive, and 3/79 buffalo and 3/86 cattle samples were *B. bigemina* positive. There was no evidence of single species infection of *B. bovis* or *B. bigemina* in any of the samples examined (Fig. 4). In the sheep samples, the prevalence of *T. lestoquardi* and *T. ovis* infections were higher rate than that of *B. ovis*. All 18 samples were *T. lestoquardi* positive, 14/18 samples were *T. ovis* positive, and 4/18 were *B. ovis* positive.

**Fig. 4.**
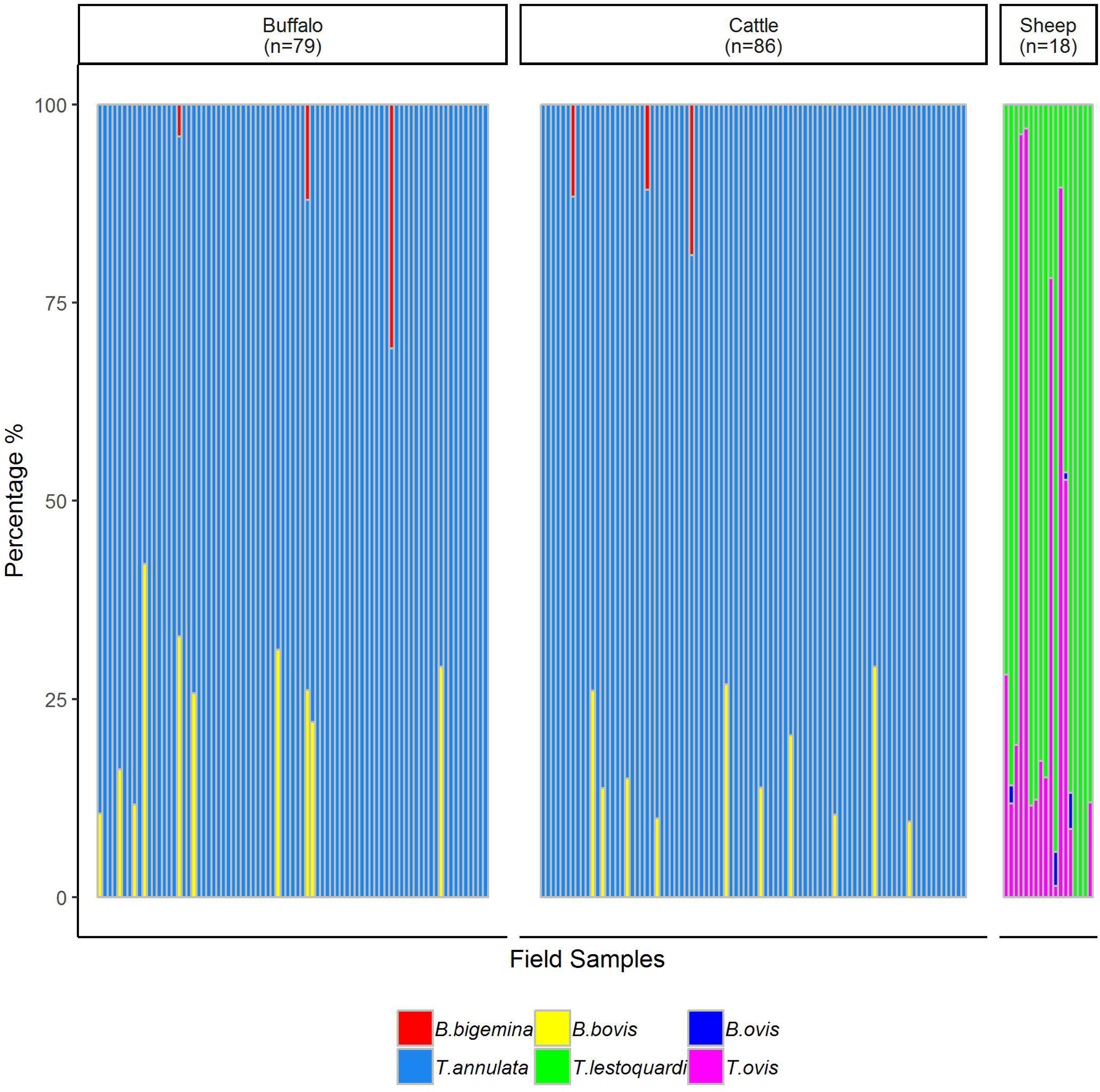
Deep amplicon sequencing for the quantification of haemoprotozoan parasites from field samples. A total of 79 positive blood samples of buffalo, 86 from cattle and 18 from sheep were collected from veterinary clinics throughout the Punjab province of Pakistan. The samples were collected into an EDTA tube and stored at −20 □C. Thick and thin blood smears were examined to identify piroplasm before extracting gDNA. The amplicon sequencing assay was applied with unknown parasitemia level of each sample. The bar of each samples shows the proportion of each species as estimated by deep amplicon sequencing assay. The Y-axis shows the percentage proportions of each species.

## Discussion

Several blood borne protozoan diseases cause important health and management problems, resulting in economic losses and reduced productivity in domestic livestock worldwide (Jabbar *et al.*, 2015). Among these, theileriosis is ranked to be most significant disease among the tick-borne pathogens of ruminants (Mans *et al.*, 2015). The most pathogenic species are *T. annulata, T. parva* and *T. lestoquardi*, while *T. mutants, T. orientalis, T. taurotragi*, and *T. ovis*, mostly cause asymptomatic infections in livestock (Nene *et al.*, 2016). Depending on the species of *Theileria*, disease is transmitted by ticks of the genera *Hyalomma, Rhipicephalus* and *Amblyomma*. The geographical distribution of *Theileria* is restricted to tropical and subtropical regions of Asia and Africa (Sivakumar *et al.*, 2014). Babesiosis infects a wide range of domestic livestock resulting in severe economic losses (Bock *et al.*, 2004). The most pathogenic species are *B. bovis, B. bigemina, B. divergens* and *B. ovis* (Sivakumar *et al.*, 2016), while *B. occultans, B. ovata, B. orientalis* and *B. major* are considered to be less pathogenic (Decaro *et al.*, 2013; He *et al.*, 2017; Ros-Garcia *et al.*, 2011; Sivakumar *et al.*, 2016). Depending on the species of *Babesia*, ticks of the genera *Boophilus* and *Ixodes* are involved in disease transmission. *Babesia* are distributed worldwide, especially in tropical and subtropical regions (Jabbar *et al.*, 2015). The severity of disease is influenced by tick infestation, climatic changes, geographical regions, co-grazing and time of the year. Haemoprotozoan species vary in terms of disease epidemiology, drug sensitivity, disease pathogenicity and control managements (Mans *et al.*, 2015).

Giemsa-stained blood smears are the standard method for the detection of haemoprotozoan in the blood of infected animals. This method is generally useful in acutely infected animals, but insensitive in the detection of chronically infected carriers, where the level of parasitaemia is low. This method is laborious and error prone in inexperienced hands (Bose *et al.*, 1995). Serological tests have been developed to detect circulatory antibodies against the parasites, but these generally have poor sensitivity and specificity due to involving cross-reactions or nonspecific immune responses (Passos *et al.*, 1998), and only detect previous exposure as opposed to current infection. Conventional PCR methods are useful in the detection of particular haemoprotozoan species for which the reagents and conditions have been developed, but can have limitations in the detection of other species and lack scalability (Bilgic *et al.*, 2013; Chaisi *et al.*, 2013; Gubbels *et al.*, 1999; Kundave *et al.*, 2018). In contrast, the deep amplicon sequencing method is potentially providing more reliable and accurate in the automated high throughput analysis of all haemoprotozoan species.

We have evaluated a deep amplicon sequencing method to determine the proportions of *Theileria* and *Babesia* species infecting large and small ruminants by: assessing sequence representation of each of the two species of *Theileria* and *Babesia* in different number of PCR cycles; validate the assay using a mock pools of each of the two species of *Theileria* and *Babesia;* and apply the method to quantify the piroplasm positive blood samples from field.

Four mock pools were made from each of the two species of *Theileria* and *Babesia* and amplified using four different numbers of PCR cycles, to assess the sequence representation of each species. We identified no significant variation in the number of first round PCR cycles in any of the four species; hence showed no sequence representation bias (Avramenko *et al.*, 2015a) arising from the number of first round PCR cycles. Nine mock pools containing different estimated proportions of each of the two species of *Theileria* and *Babesia* were generated by mixing equal amounts of gDNA. Sample series were prepared with each of the main species being absent from one mixture. The results of the mock pool were perfectly matched with no significant variations between replicates.

Accurate description of ‘haemoprotobiome’ has applications in monitoring changes in parasite diversity after drug treatment, or in understanding the impact of host immune responses, for example following vaccination against specific species, and the dynamics of piroplasm co-infections (Altay *et al.*, 2008; Nayel *et al.*, 2012). Piroplasm communities in ruminants are highly complex (Bock *et al.*, 2004; Nene *et al.*, 2016; Sivakumar *et al.*, 2014), and the potential for interspecies interaction is unexplored. Application of the deep amplicon sequencing method to describe the structure and dynamics of ‘haemoprotobiome’ could, therefore, provide a powerful solution with which to investigate co-infections in a similar manner to current trends in ‘nemobiome’(Avramenko *et al.*, 2015b; Chaudhry & Sargison, 2018) and ‘microbiome’ (Avramenko *et al.*, 2015b; Chambers, 2018; Chaudhry & Sargison, 2018; Gloor *et al.*, 2010; Rogers & Bruce, 2010) approaches that have revolutionised the study of gastrointestinal nematodes and bacteria. Having validated the Illumina Mi-Seq platform using mock pools of phenotypically verified piroplasm isolates, we applied the method to field samples collected from buffalo, cattle and sheep, throughout the Punjab province of Pakistan. This was undertaken as proof of concept to explore possibilities for the application of a high throughput practical method to determine the dynamics of piroplasm co-infections in particular *Theileria* and *Babesia* parasite species within mixed species field populations.

To summarise, we describe for the first time the use of deep amplicon sequencing using an Illumina Mi-seq platform to identify *Theileria* and *Babesia* species in mixed species populations and demonstrate its reliability on the field samples. This provides proof of concept in describing haemoprotozoan parasite communities, with potential diagnostic applications in livestock and humans in disease surveillance and epidemiology. The method can also be adapted to the study of emergence and spread of drug resistance.

## Acknowledgment

The authors of this study would like thanks Dr. Talat Naseer Pasha - Vice Chancellor of the University of Veterinary and Animal Science Lahore Pakistan for his great support in the arrangements of the sample collections.

## Supplementary Figure Legend

**S1 Fig.**
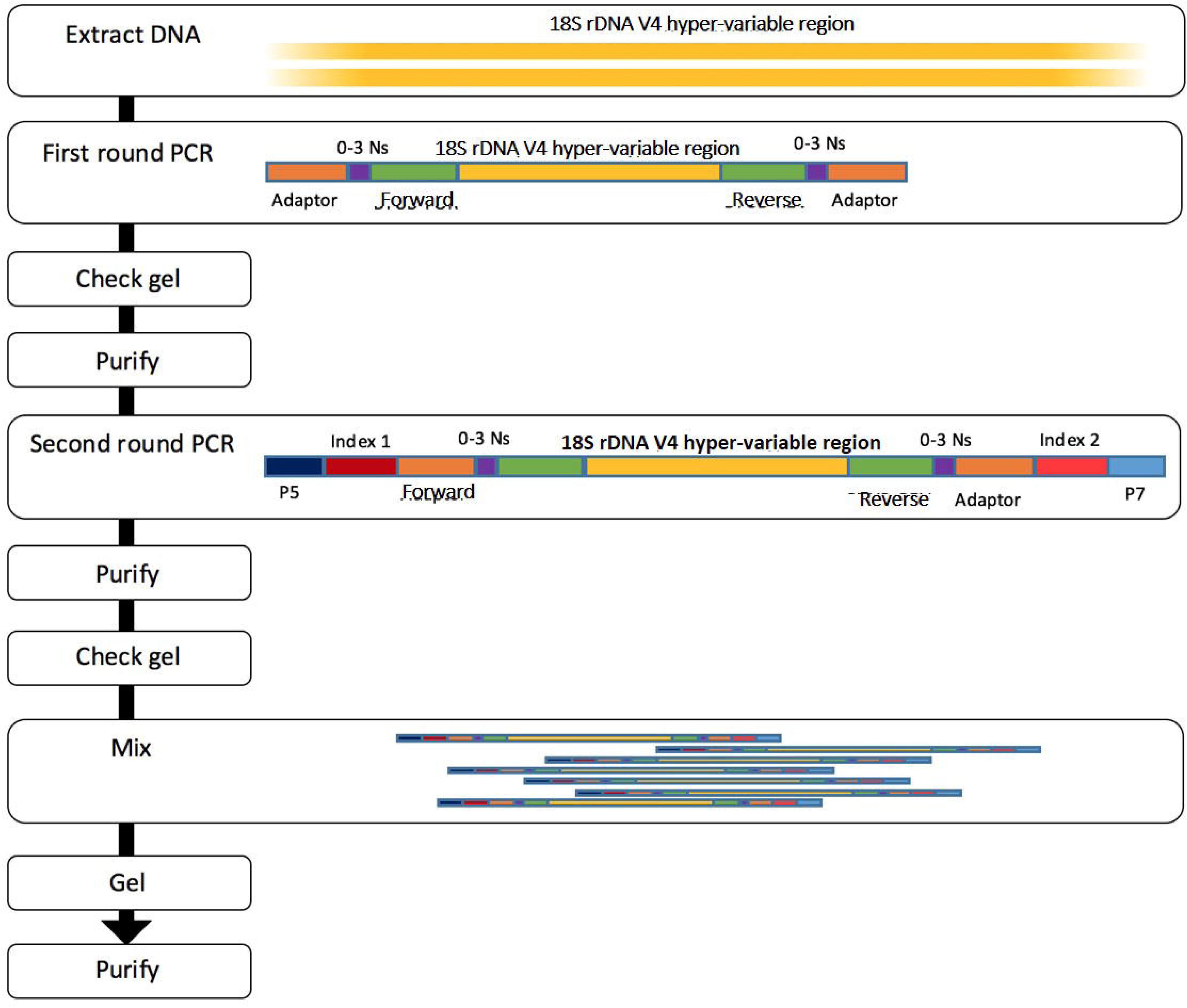
Schematic representation of the preparation of the Illumina sequencing library. In the first-round adaptor PCR amplification, overhanging primers are used to amplify the 18S rDNA to generate a fragment that includes the V4 hyper-variable region as well as the Rd1 and Rd2 specific primer regions. The Rd1 and Rd2 regions provide the target sites for the primers used for sequencing the fragment. The 0–3 random nucleotides (‘N’s) are inserted between Rd1/Rd2 sequences and the binding site of the primers to offset the reading frame, thereby increasing diversity when the amplicons are sequenced to prevent oversaturation of the MiSeq sequencing channels. A second round barcoded, limited-cycle PCR reaction is then performed using overhanging primers that bind to the Rd1 and Rd2 tags of the amplicon to add indices as well as the P5 and P7 regions required to bind to the Illumina flow cell.

