## Supplementary Table S1 for "Development of a deep amplicon sequencing method to determine the proportional species composition of piroplasm haemoprotozoa as an aid in their control"

**S1 Table. Primer sequences for Illumina MiSeq Library preparation.** RLB-For_Adp / RLB-Rev_Adp primer sequence are underlined, N’s are bolded.

| **Sequences (5'-3')** | **Primer Name** |
| --- | --- |
| TCGTCGGCAGCGTCAGATGTGTATAAGAGACAGGAGGTAGTGACAAGAAATAACAATA | RLB-For_Adp |
| TCGTCGGCAGCGTCAGATGTGTATAAGAGACAG**N**GAGGTAGTGACAAGAAATAACAATA | RLB-For_Adp1N |
| TCGTCGGCAGCGTCAGATGTGTATAAGAGACAG**NN**GAGGTAGTGACAAGAAATAACAATA | RLB-For_Adp2N |
| TCGTCGGCAGCGTCAGATGTGTATAAGAGACAG**NNN**GAGGTAGTGACAAGAAATAACAATA | RLB-For_Adp3N |
| GTCTCGTGGGCTCGGAGATGTGTATAAGAGACAGTCTTCGATCCCCTAACTTTC | RLB-Rev_Adp |
| GTCTCGTGGGCTCGGAGATGTGTATAAGAGACAG**N**TCTTCGATCCCCTAACTTTC | RLB-Rev_Adp1N |
| GTCTCGTGGGCTCGGAGATGTGTATAAGAGACAG**NN**TCTTCGATCCCCTAACTTTC | RLB-Rev_Adp2N |
| GTCTCGTGGGCTCGGAGATGTGTATAAGAGACAG**NNN**TCTTCGATCCCCTAACTTTC | RLB-Rev_dp3N |
