## Supplementary Table S2 for "Development of a deep amplicon sequencing method to determine the proportional species composition of piroplasm haemoprotozoa as an aid in their control"

**S2 Table.** **Sequences for forward and reverse barcoded primers (Nextera XT Index Kit v2).** Index sequences are highlighted. Sequences from: Oligonucleotide sequences © 2018 Illumina, Inc. All rights reserved.

| Primer | Sequence, 5’-3’ |
| --- | --- |
| S501 | AATGATACGGCGACCACCGAGATCTACACTAGATCGCTCGTCGGCAGCGTC |
| S502 | AATGATACGGCGACCACCGAGATCTACACCTCTCTATTCGTCGGCAGCGTC |
| S503 | AATGATACGGCGACCACCGAGATCTACACTATCCTCTTCGTCGGCAGCGTC |
| S504 | AATGATACGGCGACCACCGAGATCTACACAGAGTAGATCGTCGGCAGCGTC |
| S505 | AATGATACGGCGACCACCGAGATCTACACGTAAGGAGTCGTCGGCAGCGTC |
| S506 | AATGATACGGCGACCACCGAGATCTACACACTGCATATCGTCGGCAGCGTC |
| S507 | AATGATACGGCGACCACCGAGATCTACACAAGGACTATCGTCGGCAGCGTC |
| S508 | AATGATACGGCGACCACCGAGATCTACACCTAAGCCTTCGTCGGCAGCGTC |
| S510 | AATGATACGGCGACCACCGAGATCTACACCGTCTAATTCGTCGGCAGCGTC |
| S511 | AATGATACGGCGACCACCGAGATCTACACTCTCTCCGTCGTCGGCAGCGTC |
| S512 | AATGATACGGCGACCACCGAGATCTACACTCGACTAGTCGTCGGCAGCGTC |
| S513 | AATGATACGGCGACCACCGAGATCTACACTTCTAGCTTCGTCGGCAGCGTC |
| S514 | AATGATACGGCGACCACCGAGATCTACACCCTAGAGTTCGTCGGCAGCGTC |
| S515 | AATGATACGGCGACCACCGAGATCTACACGCGTAAGATCGTCGGCAGCGTC |
| S516 | AATGATACGGCGACCACCGAGATCTACACCTATTAAGTCGTCGGCAGCGTC |
| S517 | AATGATACGGCGACCACCGAGATCTACACAAGGCTATTCGTCGGCAGCGTC |
| N701 | CAAGCAGAAGACGGCATACGAGATTAAGGCGAGTCTCGTGGGCTCGG |
| N702 | CAAGCAGAAGACGGCATACGAGATCGTACTAGGTCTCGTGGGCTCGG |
| N703 | CAAGCAGAAGACGGCATACGAGATAGGCAGAAGTCTCGTGGGCTCGG |
| N704 | CAAGCAGAAGACGGCATACGAGATTCCTGAGCGTCTCGTGGGCTCGG |
| N705 | CAAGCAGAAGACGGCATACGAGATGGACTCCTGTCTCGTGGGCTCGG |
| N706 | CAAGCAGAAGACGGCATACGAGATTAGGCATGGTCTCGTGGGCTCGG |
| N707 | CAAGCAGAAGACGGCATACGAGATGTGTGTAGGTCTCGTGGGCTCGG |
| N708 | CAAGCAGAAGACGGCATACGAGATCAGAGAGGGTCTCGTGGGCTCGG |
| N709 | CAAGCAGAAGACGGCATACGAGATGCTAGGGTGTCTCGTGGGCTCGG |
| N710 | CAAGCAGAAGACGGCATACGAGATCGAGGCTGGTCTCGTGGGCTCGG |
| N711 | CAAGCAGAAGACGGCATACGAGATAAGAGGCAGTCTCGTGGGCTCGG |
| N712 | CAAGCAGAAGACGGCATACGAGATGTAGAGGAGTCTCGTGGGCTCGG |
| N713 | CAAGCAGAAGACGGCATACGAGATGCTCATGAGTCTCGTGGGCTCGG |
| N714 | CAAGCAGAAGACGGCATACGAGATATCTCAGGGTCTCGTGGGCTCGG |
| N715 | CAAGCAGAAGACGGCATACGAGATACTCGCTAGTCTCGTGGGCTCGG |
| N716 | CAAGCAGAAGACGGCATACGAGATGGAGCTACGTCTCGTGGGCTCGG |
| N717 | CAAGCAGAAGACGGCATACGAGATGCGTAGTAGTCTCGTGGGCTCGG |
| N718 | CAAGCAGAAGACGGCATACGAGATCGGAGCCTGTCTCGTGGGCTCGG |
| N719 | CAAGCAGAAGACGGCATACGAGATTACGCTGCGTCTCGTGGGCTCGG |
| N720 | CAAGCAGAAGACGGCATACGAGATATGCGCAGGTCTCGTGGGCTCGG |
| N721 | CAAGCAGAAGACGGCATACGAGATTAGCGCTCGTCTCGTGGGCTCGG |
| N722 | CAAGCAGAAGACGGCATACGAGATACTGAGCGGTCTCGTGGGCTCGG |
| N723 | CAAGCAGAAGACGGCATACGAGATCCTAAGACGTCTCGTGGGCTCGG |
| N724 | CAAGCAGAAGACGGCATACGAGATCGATCAGTGTCTCGTGGGCTCGG |
