## Supplementary Table S3 for "Development of a deep amplicon sequencing method to determine the proportional species composition of piroplasm haemoprotozoa as an aid in their control"

|  | ***T. velifera*** | ***T. taurotragi*** | ***T. sergenti*** | ***T. orientalis*** | ***T. mutans*** | ***T. buffeli*** | ***T. parva*** | ***T. annulata*** | ***B. orientalis*** | ***B. occultans*** | ***B. ovata*** | ***B. bovis*** | ***B. bigemina*** |
| --- | --- | --- | --- | --- | --- | --- | --- | --- | --- | --- | --- | --- | --- |
| ***T. taurotragi*** | 93% |  |  |  |  |  |  |  |  |  |  |  |  |
| ***T. sergenti*** | 94% | 93% |  |  |  |  |  |  |  |  |  |  |  |
| ***T. orientalis*** | 94% | 91% | 96% |  |  |  |  |  |  |  |  |  |  |
| ***T. mutans*** | 91% | 90% | 89% | 87% |  |  |  |  |  |  |  |  |  |
| ***T. buffeli*** | 94% | 93% | 98% | 98% | 89% |  |  |  |  |  |  |  |  |
| ***T. parva*** | 93% | 94% | 93% | 92% | 88% | 93% |  |  |  |  |  |  |  |
| ***T. annulata*** | 94% | 94% | 94% | 93% | 88% | 94% | 98% |  |  |  |  |  |  |
| ***B. orientalis*** | 83% | 83% | 84% | 83% | 81% | 83% | 84% | 84% |  |  |  |  |  |
| ***B. occultans*** | 84% | 83% | 85% | 84% | 82% | 84% | 86% | 85% | 97% |  |  |  |  |
| ***B. ovata*** | 82% | 82% | 82% | 81% | 82% | 81% | 83% | 83% | 92% | 92% |  |  |  |
| ***B. bovis*** | 72% | 73% | 73% | 72% | 72% | 72% | 73% | 73% | 79% | 79% | 78% |  |  |
| ***B. bigemina*** | 83% | 83% | 83% | 82% | 82% | 82% | 84% | 83% | 90% | 91% | 93% | 78% |  |
| ***B. major*** | 81% | 80% | 80% | 81% | 80% | 81% | 82% | 81% | 90% | 89% | 90% | 77% | 88% |

**S3 Table.** **Genetic distance of 18S rDNA sequences**. The data was generated from 86 rDNA 18S consensus sequences obtained from *Theleria* and *Babesia* species of large and small ruminants. Sequences were aligned with MUSCLE alignment using Geneious version 9.0.2.The distance of sequence identity (%) of each species is indicated directly each species name.

|  | ***B. ovis*** | ***T. ovis*** | ***T. lestoquardi*** | ***T. luwenshuni*** |
| --- | --- | --- | --- | --- |
| ***T. ovis*** | 82% |  |  |  |
| ***T. lestoquardi*** | 83% | 95% |  |  |
| ***T. luwenshuni*** | 81% | 95% | 93% |  |
| ***T. uilenbergi*** | 83% | 93% | 93% | 93% |
