## Supplementary Table S4 for "Development of a deep amplicon sequencing method to determine the proportional species composition of piroplasm haemoprotozoa as an aid in their control"

| **Nucleotide Position** | ***T. parva*** | ***T. annulata*** | **Nucleotide Position** | | ***T. orientalis*** | | ***T. buffeli*** | | **Nucleotide Position** | | ***T. sergenti*** | | ***T. buffeli*** | | **Nucleotide Position** | ***B. occultans*** | ***B. orientalis*** |
| --- | --- | --- | --- | --- | --- | --- | --- | --- | --- | --- | --- | --- | --- | --- | --- | --- | --- |
| 124 | A | A/G | 33 | | T | | T/C | | 33 | | T | | T/C | | 57 | T | T/C |
| 188 | C | T | 36 | | T | | T/C | | 36 | | T | | T/C | | 185 | T | T/G |
| 191 | - | T | 146 | | A/T | | A/T | | 146 | | A | | A/T | | 186 | C | T |
| 194 | T/C | T | 186 | | A | | A/T | | 186 | | A | | A/T | | 196 | T | T/C |
| 197 | C | C/T | 189 | | - | | -/A | | 188 | | T | | T/A | | 202 | G | T/G |
| 200 | T/C | T | 190 | | A/T | | A/T | | 189 | | - | | -/T | | 204 | C | G |
| 201 | T/C | C | 191 | | C | | C/T | | 190 | | T | | A/T | | 207 | A | G |
| 202 | C | T | 192 | | A | | A/T/G | | 191 | | C | | C/T | | 212 | G | A |
| 203 | G | G/T | 194 | | A/T | | A/T | | 192 | | A/T/G | | A | | 213 | T | A |
| 210 | C | G | 195 | | T | | T/A | | 194 | | A/T | | T | | 214 | C | T |
| 223 | A | T | 201 | | G | | G/T | | 195 | | T/A | | T | | 389 | C/T | T |
| 228 | G/A | G | 202 | | T | | T/- | | 201 | | G/T | | - | | 395 | G/T | G |
| 233 | G/A | G | 208 | | T | | T/G | | 202 | | T/- | | T | |  |  |  |
| 238 | C/T | T | 209 | | T/A | | T/G | | 203 | | T/C | | C | |  |  |  |
| 239 | G/A | - | 210 | | T | | T/A/C | | 208 | | T/G | | T | |  |  |  |
| 249 | G | A | 211 | | G | | G/T | | 210 | | T/A/C | | T | |  |  |  |
| 254 | T | C/T | 214 | | A/T | | A/T | | 211 | | G/T | | G | |  |  |  |
| 260 | T | T/A | 218 | | T/C | | T/C | | 214 | | T/A | | T | |  |  |  |
| 270 | A | A/G | 231 | | A/T | | - | | 218 | | T/C | | C | |  |  |  |
| 288 | T/C | T | 243 | | A | | A/T | | 231 | | A/T | | T | |  |  |  |
| 291 | T | C | 346 | | T/A | | T/A | | 321 | | A/G | | A | |  |  |  |
| 303 | T | C/T | 323 | | A/G | | A | |  |  | |  | |  |  |  |  |
| 311 | G/T | G |  |  | |  | |  |  |  |  |  |  |  |  |  |  |
| 324 | A | G/A |  |  | |  | |  |  |  |  |  |  |  |  |  |  |
| 384 | G | -/G |  |  | |  | |  |  |  |  |  |  |  |  |  |  |
| 385 | - | G |  |  | |  | |  |  |  |  |  |  |  |  |  |  |
| 478 | T | C/T |  |  | |  | |  |  |  |  |  |  |  |  |  |  |

**S4 Table.** Sequence variation in 18S rDNA of closely related species of *Theleria* and *Babesia* species.
