## Supplementary Table S5 for "Development of a deep amplicon sequencing method to determine the proportional species composition of piroplasm haemoprotozoa as an aid in their control"

| **Sample** | **Mix** | **Replicate** | ***B. bovis*** | ***B. bigemina*** | ***T. annulata*** | ***T. parva*** | **Total** |
| --- | --- | --- | --- | --- | --- | --- | --- |
| M1A | 1 | A | 25796 | 118 | 4563 | 6393 | 36870 |
| M1B | 1 | B | 23916 | 136 | 3801 | 5304 | 33157 |
| M1C | 1 | C | 17580 | 74 | 2947 | 4191 | 24792 |
| M2A | 2 | A | 20029 | 0 | 2313 | 1667 | 24009 |
| M2B | 2 | B | 21714 | 0 | 1984 | 2543 | 26241 |
| M2C | 2 | C | 22773 | 0 | 1182 | 1512 | 25467 |
| M3A | 3 | A | 0 | 495 | 7372 | 7275 | 15142 |
| M3B | 3 | B | 0 | 637 | 6657 | 7332 | 14626 |
| M3C | 3 | C | 0 | 634 | 8192 | 9695 | 18521 |
| M4A | 4 | A | 24648 | 118 | 0 | 4809 | 29575 |
| M4B | 4 | B | 25012 | 100 | 0 | 5195 | 30307 |
| M4C | 4 | C | 27491 | 51 | 0 | 4004 | 31546 |
| M5A | 5 | A | 17595 | 157 | 2469 | 0 | 20221 |
| M5B | 5 | B | 18970 | 89 | 1892 | 0 | 20951 |
| M5C | 5 | C | 19829 | 85 | 2855 | 0 | 22769 |
| M6A | 6 | A | 92756 | 0 | 0 | 0 | 92756 |
| M6B | 6 | B | 76432 | 0 | 0 | 0 | 76432 |
| M6C | 6 | C | 84563 | 0 | 0 | 0 | 84563 |
| M7A | 7 | A | 0 | 7832 | 0 | 0 | 7832 |
| M7B | 7 | B | 0 | 9432 | 0 | 0 | 9432 |
| M7C | 7 | C | 0 | 10232 | 0 | 0 | 10232 |
| M8A | 8 | A | 0 | 0 | 13858 | 0 | 13858 |
| M8B | 8 | B | 0 | 0 | 12345 | 0 | 12345 |
| M8C | 8 | C | 0 | 0 | 11467 | 0 | 11467 |
| M9A | 9 | A | 0 | 0 | 0 | 16754 | 16754 |
| M9B | 9 | B | 0 | 0 | 0 | 17564 | 17564 |
| M9C | 9 | C | 0 | 0 | 0 | 14745 | 14745 |

**S5 Table.** Two *Theileria* (*T. annulata, T. parva*) and *Babesia* species (*B. bigemina,* *B. bovis*) were used to create a variety of mock pool samples with random number of parasites from each species. The mock pools were amplified with four different numbers of cycles (25, 30, 35 and 40) to assess the accuracy and determine the species representation bias
