## Supplementary Table S7 for "Development of a deep amplicon sequencing method to determine the proportional species composition of piroplasm haemoprotozoa as an aid in their control"

**S7 Table.** Deep amplicon sequencing data of haemoprotozoan parasites from field samples. A total of 79 positive blood samples of buffalo, 86 from cattle and 18 from sheep were collected from veterinary clinics throughout the Punjab province of Pakistan.

| Host | Population | | *B. bigemina* | | *B. bovis* | | *T. annulata* | | *B. ovis* | | *T. lestoquardi* | | *T. ovis* | | total | |
| --- | --- | --- | --- | --- | --- | --- | --- | --- | --- | --- | --- | --- | --- | --- | --- | --- |
| Buffalo (n=79) | 1 | | 0 | | 5437 | | 46151 | | 0 | | 0 | | 0 | | 51588 | |
| Buffalo (n=79) | 2 | | 0 | | 0 | | 35114 | | 0 | | 0 | | 0 | | 35114 | |
| Buffalo (n=79) | 3 | | 0 | | 0 | | 60457 | | 0 | | 0 | | 0 | | 60457 | |
| Buffalo (n=79) | 4 | | 0 | | 0 | | 22743 | | 0 | | 0 | | 0 | | 22743 | |
| Buffalo (n=79) | 5 | | 0 | | 5532 | | 28758 | | 0 | | 0 | | 0 | | 34290 | |
| Buffalo (n=79) | 6 | | 0 | | 0 | | 34530 | | 0 | | 0 | | 0 | | 34530 | |
| Buffalo (n=79) | 7 | | 0 | | 0 | | 23271 | | 0 | | 0 | | 0 | | 23271 | |
| Buffalo (n=79) | 8 | | 0 | | 6534 | | 49518 | | 0 | | 0 | | 0 | | 56052 | |
| Buffalo (n=79) | 9 | | 0 | | 0 | | 43871 | | 0 | | 0 | | 0 | | 43871 | |
| Buffalo (n=79) | 10 | | 0 | | 7865 | | 10841 | | 0 | | 0 | | 0 | | 18706 | |
| Buffalo (n=79) | 11 | | 0 | | 0 | | 25541 | | 0 | | 0 | | 0 | | 25541 | |
| Buffalo (n=79) | 12 | | 0 | | 0 | | 29342 | | 0 | | 0 | | 0 | | 29342 | |
| Buffalo (n=79) | 13 | | 0 | | 0 | | 27453 | | 0 | | 0 | | 0 | | 27453 | |
| Buffalo (n=79) | 14 | | 0 | | 0 | | 38181 | | 0 | | 0 | | 0 | | 38181 | |
| Buffalo (n=79) | 15 | | 0 | | 0 | | 77261 | | 0 | | 0 | | 0 | | 77261 | |
| Buffalo (n=79) | 16 | | 0 | | 0 | | 16255 | | 0 | | 0 | | 0 | | 16255 | |
| Buffalo (n=79) | 17 | | 1073 | | 8769 | | 16814 | | 0 | | 0 | | 0 | | 26656 | |
| Buffalo (n=79) | 18 | | 0 | | 0 | | 42371 | | 0 | | 0 | | 0 | | 42371 | |
| Buffalo (n=79) | 19 | | 0 | | 0 | | 41234 | | 0 | | 0 | | 0 | | 41234 | |
| Buffalo (n=79) | 20 | | 0 | | 6785 | | 19586 | | 0 | | 0 | | 0 | | 26371 | |
| Buffalo (n=79) | 21 | | 0 | | 0 | | 19924 | | 0 | | 0 | | 0 | | 19924 | |
| Buffalo (n=79) | 22 | | 0 | | 0 | | 17718 | | 0 | | 0 | | 0 | | 17718 | |
| Buffalo (n=79) | 23 | | 0 | | 0 | | 62794 | | 0 | | 0 | | 0 | | 62794 | |
| Buffalo (n=79) | 24 | | 0 | | 0 | | 21210 | | 0 | | 0 | | 0 | | 21210 | |
| Buffalo (n=79) | 25 | | 0 | | 0 | | 12434 | | 0 | | 0 | | 0 | | 12434 | |
| Buffalo (n=79) | 26 | | 0 | | 0 | | 64120 | | 0 | | 0 | | 0 | | 64120 | |
| Buffalo (n=79) | 27 | | 0 | | 0 | | 33114 | | 0 | | 0 | | 0 | | 33114 | |
| Buffalo (n=79) | 28 | | 0 | | 0 | | 88956 | | 0 | | 0 | | 0 | | 88956 | |
| Buffalo (n=79) | 29 | | 0 | | 0 | | 56092 | | 0 | | 0 | | 0 | | 56092 | |
| Buffalo (n=79) | 30 | | 0 | | 0 | | 10218 | | 0 | | 0 | | 0 | | 10218 | |
| Buffalo (n=79) | 31 | | 0 | | 0 | | 57295 | | 0 | | 0 | | 0 | | 57295 | |
| Buffalo (n=79) | 32 | | 0 | | 0 | | 14713 | | 0 | | 0 | | 0 | | 14713 | |
| Buffalo (n=79) | 33 | | 0 | | 0 | | 10983 | | 0 | | 0 | | 0 | | 10983 | |
| Buffalo (n=79) | 34 | | 0 | | 0 | | 21325 | | 0 | | 0 | | 0 | | 21325 | |
| Buffalo (n=79) | 35 | | 0 | | 0 | | 18839 | | 0 | | 0 | | 0 | | 18839 | |
| Buffalo (n=79) | 36 | | 0 | | 0 | | 18320 | | 0 | | 0 | | 0 | | 18320 | |
| Buffalo (n=79) | 37 | | 0 | | 9765 | | 21517 | | 0 | | 0 | | 0 | | 31282 | |
| Buffalo (n=79) | 38 | | 0 | | 0 | | 21831 | | 0 | | 0 | | 0 | | 21831 | |
| Buffalo (n=79) | 39 | | 0 | | 0 | | 12993 | | 0 | | 0 | | 0 | | 12993 | |
| Buffalo (n=79) | 40 | | 0 | | 0 | | 14641 | | 0 | | 0 | | 0 | | 14641 | |
| Buffalo (n=79) | 41 | | 0 | | 0 | | 10091 | | 0 | | 0 | | 0 | | 10091 | |
| Buffalo (n=79) | 42 | | 0 | | 0 | | 12076 | | 0 | | 0 | | 0 | | 12076 | |
| Buffalo (n=79) | 43 | | 3479 | | 7564 | | 17905 | | 0 | | 0 | | 0 | | 28948 | |
| Buffalo (n=79) | 44 | | 0 | | 6689 | | 23557 | | 0 | | 0 | | 0 | | 30246 | |
| Buffalo (n=79) | 45 | | 0 | | 0 | | 17002 | | 0 | | 0 | | 0 | | 17002 | |
| Buffalo (n=79) | 46 | | 0 | | 0 | | 19012 | | 0 | | 0 | | 0 | | 19012 | |
| Buffalo (n=79) | 47 | | 0 | | 0 | | 13576 | | 0 | | 0 | | 0 | | 13576 | |
| Buffalo (n=79) | 48 | | 0 | | 0 | | 23861 | | 0 | | 0 | | 0 | | 23861 | |
| Buffalo (n=79) | 49 | | 0 | | 0 | | 14316 | | 0 | | 0 | | 0 | | 14316 | |
| Buffalo (n=79) | 50 | | 0 | | 0 | | 17344 | | 0 | | 0 | | 0 | | 17344 | |
| Buffalo (n=79) | 51 | | 0 | | 0 | | 21560 | | 0 | | 0 | | 0 | | 21560 | |
| Buffalo (n=79) | 52 | | 0 | | 0 | | 17979 | | 0 | | 0 | | 0 | | 17979 | |
| Buffalo (n=79) | 53 | | 0 | | 0 | | 12070 | | 0 | | 0 | | 0 | | 12070 | |
| Buffalo (n=79) | 54 | | 0 | | 0 | | 16959 | | 0 | | 0 | | 0 | | 16959 | |
| Buffalo (n=79) | 55 | | 0 | | 0 | | 13451 | | 0 | | 0 | | 0 | | 13451 | |
| Buffalo (n=79) | 56 | | 0 | | 0 | | 14724 | | 0 | | 0 | | 0 | | 14724 | |
| Buffalo (n=79) | 57 | | 0 | | 0 | | 16830 | | 0 | | 0 | | 0 | | 16830 | |
| Buffalo (n=79) | 58 | | 0 | | 0 | | 14491 | | 0 | | 0 | | 0 | | 14491 | |
| Buffalo (n=79) | 59 | | 0 | | 0 | | 14514 | | 0 | | 0 | | 0 | | 14514 | |
| Buffalo (n=79) | 60 | | 5437 | | 0 | | 12250 | | 0 | | 0 | | 0 | | 17687 | |
| Buffalo (n=79) | 61 | | 0 | | 0 | | 12859 | | 0 | | 0 | | 0 | | 12859 | |
| Buffalo (n=79) | 62 | | 0 | | 0 | | 16217 | | 0 | | 0 | | 0 | | 16217 | |
| Buffalo (n=79) | 63 | | 0 | | 0 | | 15116 | | 0 | | 0 | | 0 | | 15116 | |
| Buffalo (n=79) | 64 | | 0 | | 0 | | 14112 | | 0 | | 0 | | 0 | | 14112 | |
| Buffalo (n=79) | 65 | | 0 | | 0 | | 19989 | | 0 | | 0 | | 0 | | 19989 | |
| Buffalo (n=79) | 66 | | 0 | | 0 | | 14120 | | 0 | | 0 | | 0 | | 14120 | |
| Buffalo (n=79) | 67 | | 0 | | 0 | | 18184 | | 0 | | 0 | | 0 | | 18184 | |
| Buffalo (n=79) | 68 | | 0 | | 0 | | 21795 | | 0 | | 0 | | 0 | | 21795 | |
| Buffalo (n=79) | 69 | | 0 | | 0 | | 12771 | | 0 | | 0 | | 0 | | 12771 | |
| Buffalo (n=79) | 70 | | 0 | | 6456 | | 15751 | | 0 | | 0 | | 0 | | 22207 | |
| Buffalo (n=79) | 71 | | 0 | | 0 | | 12347 | | 0 | | 0 | | 0 | | 12347 | |
| Buffalo (n=79) | 72 | | 0 | | 0 | | 16264 | | 0 | | 0 | | 0 | | 16264 | |
| Buffalo (n=79) | 73 | | 0 | | 0 | | 12659 | | 0 | | 0 | | 0 | | 12659 | |
| Buffalo (n=79) | 74 | | 0 | | 0 | | 24241 | | 0 | | 0 | | 0 | | 24241 | |
| Buffalo (n=79) | 75 | | 0 | | 0 | | 21312 | | 0 | | 0 | | 0 | | 21312 | |
| Buffalo (n=79) | 76 | | 0 | | 0 | | 23923 | | 0 | | 0 | | 0 | | 23923 | |
| Buffalo (n=79) | 77 | | 0 | | 0 | | 19819 | | 0 | | 0 | | 0 | | 19819 | |
| Buffalo (n=79) | 78 | | 0 | | 0 | | 16673 | | 0 | | 0 | | 0 | | 16673 | |
| Buffalo (n=79) | 79 | | 0 | | 0 | | 13051 | | 0 | | 0 | | 0 | | 13051 | |
| Cattle (n=86) | 1 | | 0 | | 0 | | 34363 | | 0 | | 0 | | 0 | | 34363 | |
| Cattle (n=86) | 2 | | 0 | | 0 | | 23418 | | 0 | | 0 | | 0 | | 23418 | |
| Cattle (n=86) | 3 | | 0 | | 0 | | 31378 | | 0 | | 0 | | 0 | | 31378 | |
| Cattle (n=86) | 4 | | 0 | | 0 | | 28012 | | 0 | | 0 | | 0 | | 28012 | |
| Cattle (n=86) | 5 | | 0 | | 0 | | 23445 | | 0 | | 0 | | 0 | | 23445 | |
| Cattle (n=86) | 6 | | 0 | | 0 | | 45454 | | 0 | | 0 | | 0 | | 45454 | |
| Cattle (n=86) | 7 | | 4383 | | 0 | | 33251 | | 0 | | 0 | | 0 | | 37634 | |
| Cattle (n=86) | 8 | | 0 | | 0 | | 23434 | | 0 | | 0 | | 0 | | 23434 | |
| Cattle (n=86) | 9 | | 0 | | 0 | | 25123 | | 0 | | 0 | | 0 | | 25123 | |
| Cattle (n=86) | 10 | | 0 | | 0 | | 25656 | | 0 | | 0 | | 0 | | 25656 | |
| Cattle (n=86) | 11 | | 0 | | 4321 | | 12280 | | 0 | | 0 | | 0 | | 16601 | |
| Cattle (n=86) | 12 | | 0 | | 0 | | 23428 | | 0 | | 0 | | 0 | | 23428 | |
| Cattle (n=86) | 13 | | 0 | | 5432 | | 34036 | | 0 | | 0 | | 0 | | 39468 | |
| Cattle (n=86) | 14 | | 0 | | 0 | | 45321 | | 0 | | 0 | | 0 | | 45321 | |
| Cattle (n=86) | 15 | | 0 | | 0 | | 54362 | | 0 | | 0 | | 0 | | 54362 | |
| Cattle (n=86) | 16 | | 0 | | 0 | | 32414 | | 0 | | 0 | | 0 | | 32414 | |
| Cattle (n=86) | 17 | | 0 | | 0 | | 343210 | | 0 | | 0 | | 0 | | 343210 | |
| Cattle (n=86) | 18 | | 0 | | 7641 | | 43289 | | 0 | | 0 | | 0 | | 50930 | |
| Cattle (n=86) | 19 | | 0 | | 0 | | 33453 | | 0 | | 0 | | 0 | | 33453 | |
| Cattle (n=86) | 20 | | 0 | | 0 | | 28050 | | 0 | | 0 | | 0 | | 28050 | |
| Cattle (n=86) | 21 | | 0 | | 0 | | 21931 | | 0 | | 0 | | 0 | | 21931 | |
| Cattle (n=86) | 22 | | 4443 | | 0 | | 36775 | | 0 | | 0 | | 0 | | 41218 | |
| Cattle (n=86) | 23 | | 0 | | 0 | | 34014 | | 0 | | 0 | | 0 | | 34014 | |
| Cattle (n=86) | 24 | | 0 | | 5642 | | 50941 | | 0 | | 0 | | 0 | | 56583 | |
| Cattle (n=86) | 25 | | 0 | | 0 | | 17104 | | 0 | | 0 | | 0 | | 17104 | |
| Cattle (n=86) | 26 | | 0 | | 0 | | 22601 | | 0 | | 0 | | 0 | | 22601 | |
| Cattle (n=86) | 27 | | 0 | | 0 | | 49573 | | 0 | | 0 | | 0 | | 49573 | |
| Cattle (n=86) | 28 | | 0 | | 0 | | 14586 | | 0 | | 0 | | 0 | | 14586 | |
| Cattle (n=86) | 29 | | 0 | | 0 | | 16741 | | 0 | | 0 | | 0 | | 16741 | |
| Cattle (n=86) | 30 | | 0 | | 0 | | 17251 | | 0 | | 0 | | 0 | | 17251 | |
| Cattle (n=86) | 31 | | 5431 | | 0 | | 23110 | | 0 | | 0 | | 0 | | 28541 | |
| Cattle (n=86) | 32 | | 0 | | 0 | | 17830 | | 0 | | 0 | | 0 | | 17830 | |
| Cattle (n=86) | 33 | | 0 | | 0 | | 19442 | | 0 | | 0 | | 0 | | 19442 | |
| Cattle (n=86) | 34 | | 0 | | 0 | | 21314 | | 0 | | 0 | | 0 | | 21314 | |
| Cattle (n=86) | 35 | | 0 | | 0 | | 19650 | | 0 | | 0 | | 0 | | 19650 | |
| Cattle (n=86) | 36 | | 0 | | 0 | | 10651 | | 0 | | 0 | | 0 | | 10651 | |
| Cattle (n=86) | 37 | | 0 | | 0 | | 13619 | | 0 | | 0 | | 0 | | 13619 | |
| Cattle (n=86) | 38 | | 0 | | 4534 | | 12329 | | 0 | | 0 | | 0 | | 16863 | |
| Cattle (n=86) | 39 | | 0 | | 0 | | 14940 | | 0 | | 0 | | 0 | | 14940 | |
| Cattle (n=86) | 40 | | 0 | | 0 | | 13519 | | 0 | | 0 | | 0 | | 13519 | |
| Cattle (n=86) | 41 | | 0 | | 0 | | 22108 | | 0 | | 0 | | 0 | | 22108 | |
| Cattle (n=86) | 42 | | 0 | | 0 | | 21026 | | 0 | | 0 | | 0 | | 21026 | |
| Cattle (n=86) | 43 | | 0 | | 0 | | 13046 | | 0 | | 0 | | 0 | | 13046 | |
| Cattle (n=86) | 44 | | 0 | | 0 | | 12605 | | 0 | | 0 | | 0 | | 12605 | |
| Cattle (n=86) | 45 | | 0 | | 5431 | | 33766 | | 0 | | 0 | | 0 | | 39197 | |
| Cattle (n=86) | 46 | | 0 | | 0 | | 15539 | | 0 | | 0 | | 0 | | 15539 | |
| Cattle (n=86) | 47 | | 0 | | 0 | | 12132 | | 0 | | 0 | | 0 | | 12132 | |
| Cattle (n=86) | 48 | | 0 | | 0 | | 11157 | | 0 | | 0 | | 0 | | 11157 | |
| Cattle (n=86) | 49 | | 0 | | 0 | | 23414 | | 0 | | 0 | | 0 | | 23414 | |
| Cattle (n=86) | 50 | | 0 | | 0 | | 18606 | | 0 | | 0 | | 0 | | 18606 | |
| Cattle (n=86) | 51 | | 0 | | 6542 | | 25476 | | 0 | | 0 | | 0 | | 32018 | |
| Cattle (n=86) | 52 | | 0 | | 0 | | 13959 | | 0 | | 0 | | 0 | | 13959 | |
| Cattle (n=86) | 53 | | 0 | | 0 | | 14149 | | 0 | | 0 | | 0 | | 14149 | |
| Cattle (n=86) | 54 | | 0 | | 0 | | 34200 | | 0 | | 0 | | 0 | | 34200 | |
| Cattle (n=86) | 55 | | 0 | | 0 | | 21786 | | 0 | | 0 | | 0 | | 21786 | |
| Cattle (n=86) | 56 | | 0 | | 0 | | 16152 | | 0 | | 0 | | 0 | | 16152 | |
| Cattle (n=86) | 57 | | 0 | | 0 | | 16544 | | 0 | | 0 | | 0 | | 16544 | |
| Cattle (n=86) | 58 | | 0 | | 0 | | 18699 | | 0 | | 0 | | 0 | | 18699 | |
| Cattle (n=86) | 59 | | 0 | | 0 | | 14427 | | 0 | | 0 | | 0 | | 14427 | |
| Cattle (n=86) | 60 | | 0 | | 3573 | | 30746 | | 0 | | 0 | | 0 | | 34319 | |
| Cattle (n=86) | 61 | | 0 | | 0 | | 13236 | | 0 | | 0 | | 0 | | 13236 | |
| Cattle (n=86) | 62 | | 0 | | 0 | | 18760 | | 0 | | 0 | | 0 | | 18760 | |
| Cattle (n=86) | 63 | | 0 | | 0 | | 13397 | | 0 | | 0 | | 0 | | 13397 | |
| Cattle (n=86) | 64 | | 0 | | 0 | | 21620 | | 0 | | 0 | | 0 | | 21620 | |
| Cattle (n=86) | 65 | | 0 | | 0 | | 18343 | | 0 | | 0 | | 0 | | 18343 | |
| Cattle (n=86) | 66 | | 0 | | 0 | | 14279 | | 0 | | 0 | | 0 | | 14279 | |
| Cattle (n=86) | 67 | | 0 | | 0 | | 24548 | | 0 | | 0 | | 0 | | 24548 | |
| Cattle (n=86) | 68 | | 0 | | 5431 | | 13230 | | 0 | | 0 | | 0 | | 18661 | |
| Cattle (n=86) | 69 | | 0 | | 0 | | 26251 | | 0 | | 0 | | 0 | | 26251 | |
| Cattle (n=86) | 70 | | 0 | | 0 | | 10033 | | 0 | | 0 | | 0 | | 10033 | |
| Cattle (n=86) | 71 | | 0 | | 0 | | 11459 | | 0 | | 0 | | 0 | | 11459 | |
| Cattle (n=86) | 72 | | 0 | | 0 | | 4559 | | 0 | | 0 | | 0 | | 4559 | |
| Cattle (n=86) | 73 | | 0 | | 0 | | 12436 | | 0 | | 0 | | 0 | | 12436 | |
| Cattle (n=86) | 74 | | 0 | | 0 | | 20099 | | 0 | | 0 | | 0 | | 20099 | |
| Cattle (n=86) | 75 | | 0 | | 3422 | | 32410 | | 0 | | 0 | | 0 | | 35832 | |
| Cattle (n=86) | 76 | | 0 | | 0 | | 15076 | | 0 | | 0 | | 0 | | 15076 | |
| Cattle (n=86) | 77 | | 0 | | 0 | | 21417 | | 0 | | 0 | | 0 | | 21417 | |
| Cattle (n=86) | 78 | | 0 | | 0 | | 11241 | | 0 | | 0 | | 0 | | 11241 | |
| Cattle (n=86) | 79 | | 0 | | 0 | | 14995 | | 0 | | 0 | | 0 | | 14995 | |
| Cattle (n=86) | 80 | | 0 | | 0 | | 13412 | | 0 | | 0 | | 0 | | 13412 | |
| Cattle (n=86) | 81 | | 0 | | 0 | | 22353 | | 0 | | 0 | | 0 | | 22353 | |
| Cattle (n=86) | 82 | | 0 | | 0 | | 10100 | | 0 | | 0 | | 0 | | 10100 | |
| Cattle (n=86) | 83 | | 0 | | 0 | | 13457 | | 0 | | 0 | | 0 | | 13457 | |
| Cattle (n=86) | 84 | | 0 | | 0 | | 16785 | | 0 | | 0 | | 0 | | 16785 | |
| Cattle (n=86) | 85 | | 0 | | 0 | | 13800 | | 0 | | 0 | | 0 | | 13800 | |
| Cattle (n=86) | 86 | | 0 | | 0 | | 13430 | | 0 | | 0 | | 0 | | 13430 | |
| Sheep (n=18) | 1 | | 0 | | 0 | | 0 | | 0 | | 10588 | | 4139 | | 14727 | |
| Sheep (n=18) | 2 | | 0 | | 0 | | 0 | | 457 | | 16962 | | 2331 | | 19750 | |
| Sheep (n=18) | 3 | | 0 | | 0 | | 0 | | 0 | | 11543 | | 2741 | | 14284 | |
| Sheep (n=18) | 4 | | 0 | | 0 | | 0 | | 0 | | 1373 | | 35025 | | 36398 | |
| Sheep (n=18) | 5 | | 0 | | 0 | | 0 | | 0 | | 1225 | | 38821 | | 40046 | |
| Sheep (n=18) | 6 | | 0 | | 0 | | 0 | | 0 | | 14196 | | 1854 | | 16050 | |
| Sheep (n=18) | 7 | | 0 | | 0 | | 0 | | 0 | | 12353 | | 1728 | | 14081 | |
| Sheep (n=18) | 8 | | 0 | | 0 | | 0 | | 0 | | 10034 | | 2079 | | 12113 | |
| Sheep (n=18) | 9 | | 0 | | 0 | | 0 | | 0 | | 17825 | | 3178 | | 21003 | |
| Sheep (n=18) | 10 | | 0 | | 0 | | 0 | | 0 | | 10605 | | 37819 | | 48424 | |
| Sheep (n=18) | 11 | | 0 | | 0 | | 0 | | 1372 | | 30170 | | 456 | | 31998 | |
| Sheep (n=18) | 12 | | 0 | | 0 | | 0 | | 0 | | 1542 | | 13217 | | 14759 | |
| Sheep (n=18) | 13 | | 0 | | 0 | | 0 | | 1123 | | 53666 | | 60916 | | 115705 | |
| Sheep (n=18) | 14 | | 0 | | 0 | | 0 | | 762 | | 14510 | | 1439 | | 16711 | |
| Sheep (n=18) | | 15 | | 0 | | 0 | | 0 | | 0 | | 18711 | | 0 | | 18711 |
| Sheep (n=18) | | 16 | | 0 | | 0 | | 0 | | 0 | | 11403 | | 0 | | 11403 |
| Sheep (n=18) | | 17 | | 0 | | 0 | | 0 | | 0 | | 1890 | | 0 | | 1890 |
| Sheep (n=18) | | 18 | | 0 | | 0 | | 0 | | 0 | | 24263 | | 3303 | | 27566 |
